# Fast Connectivity Gradient Approximation: Maintaining spatially fine-grained connectivity gradients while reducing computational costs

**DOI:** 10.1101/2023.07.22.550017

**Authors:** Karl-Heinz Nenning, Ting Xu, Arielle Tambini, Alexandre R. Franco, Daniel S. Margulies, Stanley J. Colcombe, Michael P. Milham

## Abstract

Brain connectome analysis suffers from the high dimensionality of connectivity data, often forcing a reduced representation of the brain at a lower spatial resolution or parcellation. However, maintaining high spatial resolution can both allow fine-grained topographical analysis and preserve subtle individual differences otherwise lost. This work presents a computationally efficient approach to estimate spatially fine-grained connectivity gradients and demonstrates its application in improving brain-behavior predictions.

## Main

Graph-based models have become mainstream tools for the visualization and characterization of relationship structures within complex systems in machine learning, computer vision, and bioinformatics. Unfortunately, their application is not without challenges - particularly when dealing with representations at higher resolutions, as their computational and storage demands can exceed the capacity of most computing resources. Data reduction is indispensable when the feasibility of graph-based models is impacted. Such reductions can be achieved through statistical techniques (e.g., principal component, factor analyses), data downsampling approaches (e.g., local averaging based on spatial or topographic structure), or both. Here, using graph-based models in neuroimaging as an example, we demonstrate the need for careful consideration of the order in which these two reduction strategies are applied, suggesting an optimized workflow when both are pursued.

In neuroimaging, graph-based representations are increasingly used to characterize topographic patterns of gradual transitions and boundaries between systems - i.e., connectivity gradients.^1–3^ But maintaining a full spatial representation of the connectome is challenging. The number of connections scales exponentially with the number of nodes, and a common spatial resolution of the brain comprising 60,000 cortical nodes (vertices) constitutes 3,600,000,000 connections (edges). This makes it virtually impossible to evaluate the full-scale connectivity structure on consumer grade hardware. A common practice to overcome this challenge is to reduce the spatial resolution of the data, typically by averaging across vertices within spatial regions of interest (ROIs) defined using a parcellation template.^4^ However, the loss of individual-specific detail in such parcellation approaches may result in the loss of meaningful information; there is also little agreement on the appropriate choice of parcellation template in connectome studies.^5,6^ Moreover, the capability to maintain a fine-grained spatial resolution allows researchers to fully characterize the spatial layout of functional regions, an important and often overlooked aspect of modeling brain connectivity.^7^

Here, to facilitate connectivity gradients at a fine-grained spatial resolution, we propose *Fast Connectivity Gradient Approximation* (FCGA). Instead of using a full-scale vertex-by-vertex connectivity matrix, at its core, FCGA leverages a set of landmarks to approximate the underlying connectivity structure at full spatial resolution, with an efficiency that makes it possible to run on common computer hardware. These landmarks are flexible, and can, for example, be based on individual vertices or predefined ROIs.

A schematic workflow of our proposed approach is displayed in Figure 1a. After (i) establishing a set of k landmarks, the (ii) functional connectivity between all n vertices (or voxels) and the k landmarks is calculated, resulting in a n-by-k connectivity matrix (CM_n-by-k_) between all pairs of vertices and landmarks. Following current practices when calculating connectivity gradients, we quantify the spatial similarity of vertex-wise connectivity profiles. To do so, (iii) a k-by-k (landmark-by-landmark) connectivity matrix (CM_k-by-k_) that characterizes the functional connectivity between all pairs of landmarks is established. Then, row-wise thresholding, retaining the 10% strongest positive connections, is applied to both CM_n-by-k_ and CM_k-by-k_. Subsequently, (iv) the row-wise cosine similarity between all pairs of vertices in CM_n-by-k_ and CM_k-by-k_ is calculated. This results in an n-by-k affinity matrix (W_n-by-k_) that characterizes the spatial similarities between the voxel-wise connectivity profiles with the landmark regions. Finally, (v) we use Principal Component Analysis for dimensionality reduction of W_n-by-k_, establishing the approximated connectivity gradients.

**Figure 1.**
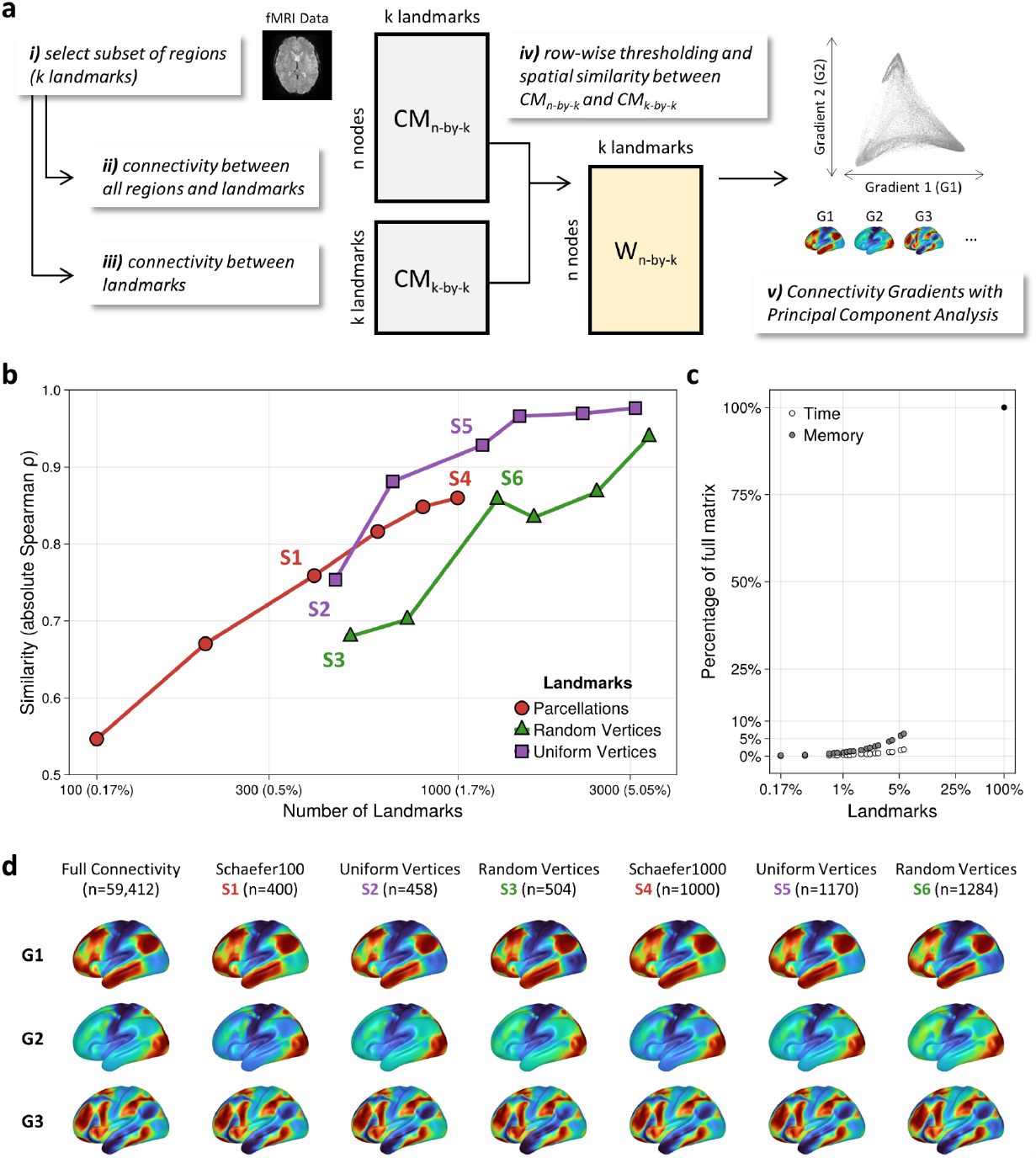
a) A schematic workflow of the proposed Fast Connectivity Gradient Approximation (FCGA) approach. b) FCGA provides high fidelity to the full connectivity structure with only a fraction of landmarks and c) <10% computational time and memory requirements. d) The core topography is similar across a different number of landmarks and sampling choices.

We evaluated the validity of FCGA on the group and individual level, using high resolution fMRI data (59,412 cortical vertices) from the Human Connectome Project (HCP)^8^ and the Nathan Kline Institute-Rockland Sample (NKI-RS).^9^ We parametrically varied the amount of downsampling (number of landmarks) across sampling choices (vertex- and ROI-level).

At the group level, we used Spearman rank correlation to compare the spatial similarity between the approximated connectivity gradients (G_FCGA_) and the connectivity gradients based on the full 59,412-by-59,412 connectivity matrix (G_full_fc_). The spatial similarity between G_FCGA_ and G_full_fc_ increased with the number of landmarks used to calculate the approximated gradients (Figure 1b). Remarkably, using 1000 ROIs as landmarks (∼1.7% of the full connectivity matrix), the average spatial similarity across 25 connectivity gradients reached ρ=0.86. Using 3000 (∼5%) uniformly distributed vertices, a spatial similarity of ρ=0.98 was achieved with <10% of the computational time and memory usage as compared to the calculation of G_full_fc_ (Figure 1c). The spatial topography of the top connectivity gradients, shown in Figure 1d, further illustrates the high spatial similarity between G_FCGA_ and G_full_fc_ across a number of sampling choices.

At the individual level, reliability and discriminability analysis confirmed the repeatability and the preservation of individual features with G_FCGA_. To quantify agreement between G_FCGA_ and G_full_fc_, we calculated the intraclass correlation coefficient (ICC) and discriminability for the same acquisition. We observed an average vertex-wise ICC of >0.9 from 1000 (∼1.7%) landmarks onwards, and discriminability was already nearly perfect with 300 (<1%) landmarks (Figure 2a). Next, we compared the ICC and the discriminability of G_FCGA_ and G_full_fc_ across sessions (Figure 2b). We observed that landmarks based on single vertices (uniformly or randomly distributed) yielded slightly lower ICC and discriminability for G_FCGA_ than for G_full_fc_. However, landmarks based on predefined parcels on the group or individual level yielded a slightly better discriminability and a similar ICC for G_FCGA_ when compared to G_full_fc_ (Figure 2b).

**Figure 2.**
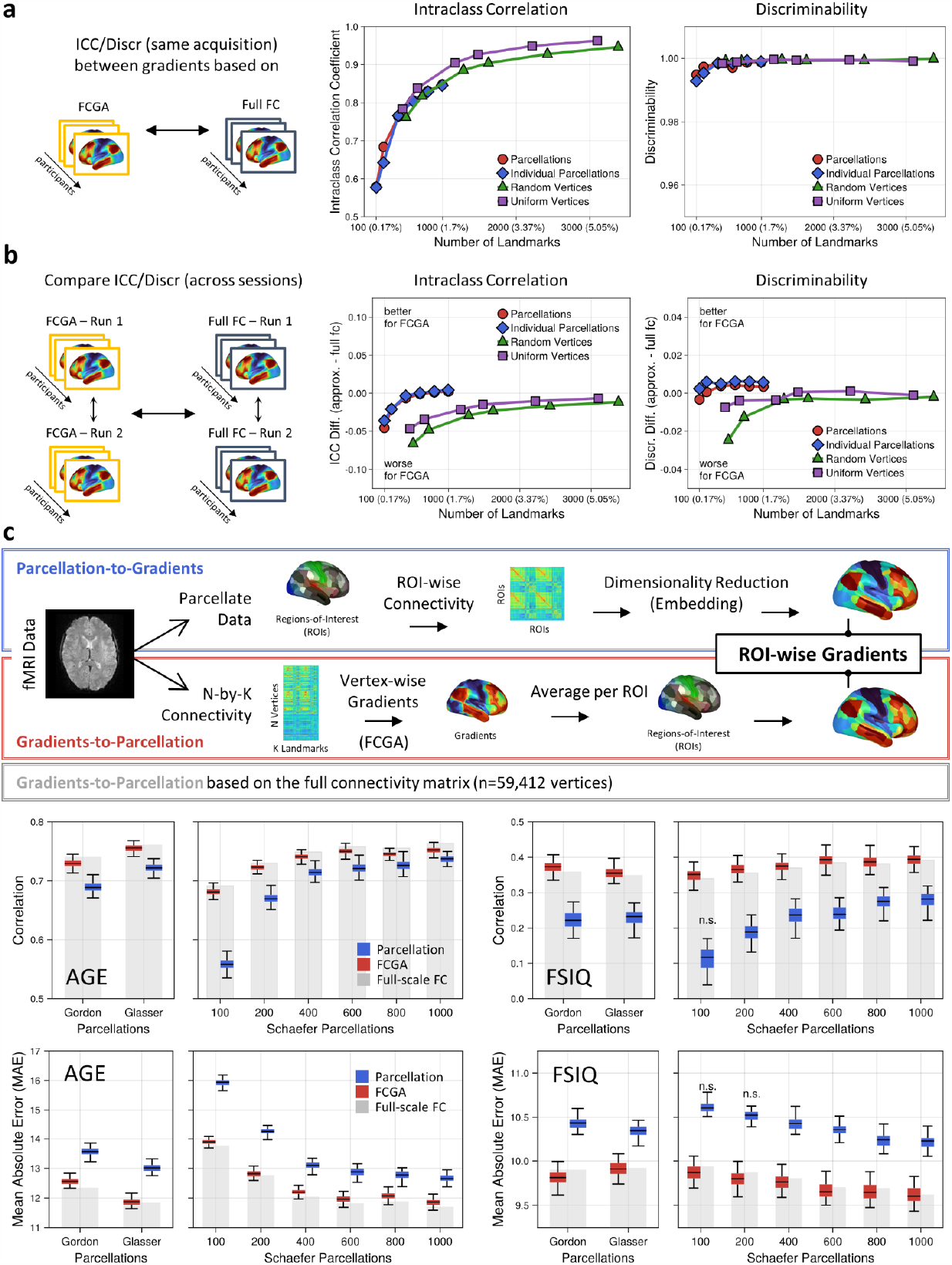
Intraclass correlation and discriminability within the same session confirms the repeatability and the preservation of individual features within (a) and across (b) sessions. c) Parcel-averaging of spatially fine-grained gradients outperforms gradient calculation on a parcel level for both age and FSIQ prediction. Boxplot show 500 predictions, colored boxes interquartile range (iqr) with whiskers spanning 1.5*iqr. Predictions that are not significantly better than prediction with randomly shuffled labels are denoted as n.s.

Finally, we evaluated the practical implications of our FCGA approach on brain-wide association studies by predicting age and full-scale intelligence (FSIQ) across the lifespan. Specifically, we compared the predictive performance of coarse-grained gradients calculated from parcellated time-series to FCGA constructed parcel-wise summarized fine-grained gradients (Figure 2c). Importantly, we observed that fine-grained gradients (i.e. G_FCGA_) subjected to parcel-averaging outperformed coarse-grained gradients for predicting both age and FSIQ across all tested parcellations (p_Bonferroni_<0.05 in 32/32 comparisons) (Figure 2c). Notably, for FSIQ we observed that gradients calculated on a low number of parcels did not perform better than chance. This indicates that constructing connectivity gradients from parcellated data might lose meaningful information in the process that is otherwise preserved in the fine-grained gradient topography.

Taken together, our results demonstrate the feasibility of establishing spatially fine-grained connectivity gradients while mitigating the computational burden of vertex-level functional connectivity data by reducing computational time and memory usage. The ability to fully appreciate the spatial layout of functional regions in gradient representations at full resolution could improve associations between functional imaging data and cognitive traits,^7^ and inform structure-function relationships.^10^ Importantly, the advantage of preserving informative individual features in gradients of spatially fine-grained connectivity data was emphasized by the improved prediction of age and intelligence over gradients based on coarse-grained connectivity data. Overall, FCGA strengthens the core concept of connectivity gradients by maintaining the full spatial resolution to study spatial transitions of brain organization. Finally, its efficiency also removes computational barriers and can enable the widespread application of connectivity gradients to capture signatures of the connectome.

## Methods

### Data

To evaluate the validity of FCGA on the group and individual level, we used resting-state fMRI data from the Human Connectome Project (HCP).^8^ The HCP data was acquired at Washington University at St. Louis on a customized Siemens 3T Connectome Skyra scanner. Resting-state fMRI was acquired with a multiband factor of 8, 2mm isotropic resolution, and a repetition time of 0.72 seconds for a duration of 14.4 minutes, resulting in 1200 volumes per acquisition. Participants were asked to relax, keep eyes open and fixated on a crosshair, and not to fall asleep. Four resting-state runs were collected in different sessions across two days (REST1 and REST2), where each session comprised two runs with different phase encoding directions (LR and RL). We used the minimally preprocessed fMRI data provided by the HCP.^11^ Briefly, the resting-state data has been motion corrected, minimally spatially smoothed (2mm), high-pass filtered (2000s cutoff), denoised for motion-related confounds and artifacts using independent component analysis,^12^ and spatially aligned to the 2 mm standard CIFTI grayordinates space.^11,13^

At the group level, we used the dense group-average functional connectome provided as part of the HCP S1200 data release. In brief, this connectivity matrix was constructed from the minimal preprocessed data across 1003 individuals that each had ∼1 hour (4x14.4 minutes) of resting-state fMRI acquisitions. The dense connectivity matrix of size 91,282 x 91,282 grayordinates (59,412 cortical vertices and 31,870 subcortical voxels) was calculated based on components of an incremental group-pca. For the group-average connectome of the HCP, we averaged the precomputed functional connectivity measures for each landmark instead of recalculating landmark-based connectivity. At individual-level, we used the 100 unrelated subjects cohort from the HCP (54F/46M, age = 29 ± 3.7 years) and two repeated resting-state fMRI runs (REST1_LR and REST2_LR) for each individual. We used the minimally preprocessed data as provided by the HCP. For both the group and individual level, we focused our analysis on the 59,412 cortical vertices.

We used a cross-sectional lifespan sample with 313 healthy participants (214 female, age: 6-85 years, 42.2 ± 22.4 years) to evaluate the practical implications of our approach on brain-wide association studies. Participants were selected from the Nathan Kline Institute-Rockland Sample (NKI-RS) ^9^, who have no diagnosis of any mental or neurological disorders, and passed quality control of a head motion criteria (mean framewise displacement < 0.25mm). The NKI-RS data was acquired at the Nathan Kline Institute on a Siemens TrioTim 3 Tesla scanner. Resting-state fMRI was acquired with a multiband factor of 4, 3mm isotropic resolution, and a repetition time of 0.645 seconds for a duration of 9.7 minutes, which resulted in 900 volumes per run. Preprocessing was performed with the Connectome Computational System,^14^ and included discarding the first five timepoints, compressing temporal spikes, slice timing correction, motion correction, 4D global mean intensity normalization, nuisance regression (Friston’s 24 model, cerebrospinal fluid and white matter), linear and quadratic detrending, band-pass filtering (0.01-0.1Hz), as well as global signal regression. The preprocessed data were then projected on the 32k fsLR surface template with 32,492 vertices per hemisphere.

### Connectivity landmarks

We used a varying number of landmarks across different sampling strategies. We utilized randomly and uniformly distributed vertices across the cortex, where the selected vertices were consistent across individuals. On the ROI-level, connectivity landmarks were defined by the average time-series based on group parcellations^15–17^ and individualized parcellations for the HCP sample.^18^

### Comparing gradients

For the group level analysis and comparisons, no alignment or reordering of the gradients (PCA components) was performed. For the analysis on the individual level, we used one set of reference gradients based on the HCP dense connectome for both the approximated and full-scale gradients. For each individual, sampling, and gradient construction approaches, the gradients were aligned to this reference with orthogonal Procrustes alignment.^19^ Orthogonal Procrustes finds the optimal linear transformation so that two sets of gradients have a matched component order and coefficient signs.

### Intraclass correlation and discriminability

We compared the reliability and discriminability of the approximated gradients (G_FCGA_) and the full-scale gradients (G_full_fc_) within and across sessions, using 2 repeated resting-state acquisitions from the 100 unrelated individuals sample of the HCP. Within-session analysis treated G_FCGA_ and G_full_fc_ of the same acquisition as repeated measures. Across-session analysis evaluated differences in the reliability and reproducibility between G_FCGA_ and G_full_fc_. Discriminability^20^ was used to measure the similarity of connectivity gradients. It is a nonparametric multivariate statistic that quantifies the degree to which repeated measurements (e.g., gradients of different approaches or sessions) are relatively similar to each other. At the vertex level, we quantified the intraclass correlation coefficient (ICC), a univariate measure of the degree of absolute agreement.^21,22^

### Prediction of individual-specific measures

We used the NKI-RS lifespan sample to evaluate implications of our proposed approach for the prediction of individual-specific measures such as age and FSIQ. As described above, connectivity gradients are often calculated on a reduced data representation on a parcel-level. In this study, we compared the predictive performance of connectivity gradients calculated on parcellated time-series (parcellation-to-gradients) to a parcel-wise averaging of fine-scale gradients constructed with FCGA (gradients-to-parcellation). Connectivity gradients of the parcellated data are established as in prior work,^1^ using all parcels as landmarks instead of a subset. We evaluated distinct, commonly used parcellations,^15–17^ and focused on the first 5 gradients as features to avoid excessively increasing the feature space. For prediction, we used ridge regression with a L2 regularization as implemented with glmnet,^23^ and a nested 10-fold cross-validation scheme for hyperparameter (lambda) selection. The predictions of each fold were aggregated, and performance was measured with mean absolute error (MAE) and correlation to the true age and FSIQ values. We repeated the 10-fold cross-validation runs 500 times, with random splits for each fold. Additionally, we tested the prediction results against a baseline of 500 prediction runs with randomly shuffled labels to evaluate if the predictive performance is greater than chance.

